# PARKIN REGULATES DRUG TAKING-BEHAVIOR IN RAT MODEL OF METHAMPHETAMINE USE DISORDER

**DOI:** 10.1101/2020.11.24.394825

**Authors:** Akhil Sharma, Arman Harutyunyan, Bernard L. Schneider, Anna Moszczynska

**Author notes:** Correspondence should be addressed to: Anna Moszczynska, Ph.D., Department of Pharmaceutical Sciences, Eugene Applebaum College of Pharmacy and Health Sciences, Wayne State University, 259 Mack Ave, Detroit, MI, USA 48201.

## Abstract

There is no FDA-approved medication for Methamphetamine (METH) Use Disorder. New therapeutic approaches are needed, especially for people who use METH heavily and are at high risk for overdose. This study used genetically engineered rats to evaluate PARKIN as a potential target for METH Use Disorder. PARKIN knockout, PARKIN-overexpressing and wild-type young adult male Long Evans rats were trained to self-administer high doses of METH using an extended-access METH self-administration paradigm. Reinforcing/rewarding properties of METH were assessed by quantifying drug-taking behavior and time spent in a METH-paired environment. PARKIN knockout rats self-administered more METH and spent more time in the METH-paired environment than wild-type rats. Wild-type rats overexpressing PARKIN self-administered less METH and spent less time in the METH-paired environment. PARKIN knockout rats overexpressing PARKIN self-administered less METH during the first half of drug self-administration days than PARKIN-deficient rats. The results indicate that rats with PARKIN excess or PARKIN deficit are useful models for studying neural substrates underlying “resilience” or vulnerability to METH Use Disorder, and identify PARKIN as a novel potential drug target to treat heavy use of METH.

## INTRODUCTION

Methamphetamine (METH) Use Disorder is a world-wide health problem. In the United States, there are more than 900,000 people who use METH and deaths from METH overdose are rapidly rising^1,2^. Despite numerous clinical trials conducted to date, there is no FDA-approved medication for METH Use Disorder. The medications tested in clinical trials have shown low efficacy in people who use METH moderately and no effect in those who use METH heavily^3–5^. New therapeutic approaches are needed, particularly for people who use METH heavily, as they suffer the most from METH abuse-related neuropsychological problems^6–8^, are less likely to seek treatment than those using the drug moderately^9^, and are at high risk of dying from METH overdose^10^. Early intervention in METH abuse by lowering METH intake is essential not only for preventing METH overdose, but also for subsequent interventions, as greater treatment participation is achieved when METH use is low^9^.

METH Use Disorder has been linked to alterations in dopaminergic and glutamatergic neurotransmission, alterations in energy metabolism and cytoskeletal arrangement as well as to oxidative stress and inflammation^11–24^. Protein-ubiquitin ligase PARKIN may be a potential novel drug candidate in METH Use Disorder as its function has been linked to these processes^25–31^. Despite evidence for a potential role of PARKIN in METH Use Disorder, ours is the first study linking PARKIN to this disorder.

Drug addiction is a chronic, relapsing disorder that has been characterized by compulsive seeking and escalated intake of drugs. In the current study, we used genetically engineered rats to evaluate PARKIN as a potential target for METH Use Disorder. PARKIN knockout (PKO), PARKIN-overexpressing (PO) and wild-type (WT) young adult male Long Evans rats were trained to self-administer high doses of METH using an extended-access METH self-administration (EA METH SA) paradigm with three progressive ratio schedules. This paradigm produced an escalation of METH intake with rats working much harder to obtain METH with increasing ratio schedule and, therefore, modeled an important aspect of human drug abuse (compulsive drug taking)^32^ and heavy consumption of METH seen in some people dependent on METH. Binging observed in people who use METH heavily can last 3-15 days. To approximate binging behavior, rats were allowed to self-administer high METH doses for 10 days. Rewarding properties of METH were assessed by measuring the number of lever presses for METH and METH intake during each of 10 operant sessions. Rewarding properties of METH were assessed by measuring time spent in a METH-paired environment. We demonstrate that PKO rats are predisposed to heavy METH use compared to WT counterparts whereas rats overexpressing PARKIN in the nucleus accumbens (a key area mediating rewarding/reinforcing effects) are less predisposed.

## MATERIALS AND METHODS

### Animals

The study employed young adult male Long Evans rats (~55 days-old at the beginning of the study, N=177) of four genotypes: WT rats, PKO rats (*Park2*^−/−^), PO rats, and PKO PO rats (Horizon Discovery, Missouri, MO/Envigo, Indianapolis, IN, USA). PO rats were generated by bilaterally overexpressing PARKIN in the nucleus accumbens of WT rats whereas PKO PO rats were generated by bilaterally overexpressing PARKIN in the nucleus accumbens of PKO rats. The stereotaxic surgery was performed as previously published33, with some modifications. Microinjections were done at a 16° angle into these coordinates: +1.8 A/P, ±3.2.M/L, −7.6 D/V from bregma. Validation of PARKIN protein loss in PKO rats is shown in Fig.2 whereas validation of parkin overexpression is shown in Fig.3. Upon arrival, animals were pair-housed and maintained under standard environmental conditions in an AAALAC-accredited vivarium. Animals were maintained on a 12 h light/dark cycle with *ad libitum* access to food and water unless specified otherwise. Body weight was monitored on a daily basis before and during experiments (Suppl. Fig.S1). All experiments were approved by the Wayne State University Institutional Animal Care and Use Committee.

### Extended-Access Methamphetamine Self-Administration (EA METH SA) and Conditioned Place Preference Test

In the first experiment, PKO and WT rats were trained to self-administer high doses of METH (measurement of METH reinforcement) (Fig.1A) or subjected to the conditioned place preference test (measurement of METH reward) (Fig.2A). In the second experiment, the reinforcing/rewarding properties of METH were assessed in PO rats as compared to WT rats. In the third experiment, PARKIN was overexpressed in the nucleus accumbens of PKO rats that were subsequently compared to non-overexpressing PKO rats (phenotype rescue experiment). During EA METH SA, rats had access to METH (0.1mg/kg/injection) for 15h/d for 10d at increasing fixed ratio (FR): FR1 (days 1-3), FR2 (days 4-6) and FR5 (days 7-10). During METH conditioning for conditioned place preference test, rats were injected with 4mg/kg METH or saline on alternate days.

**Figure 1.**
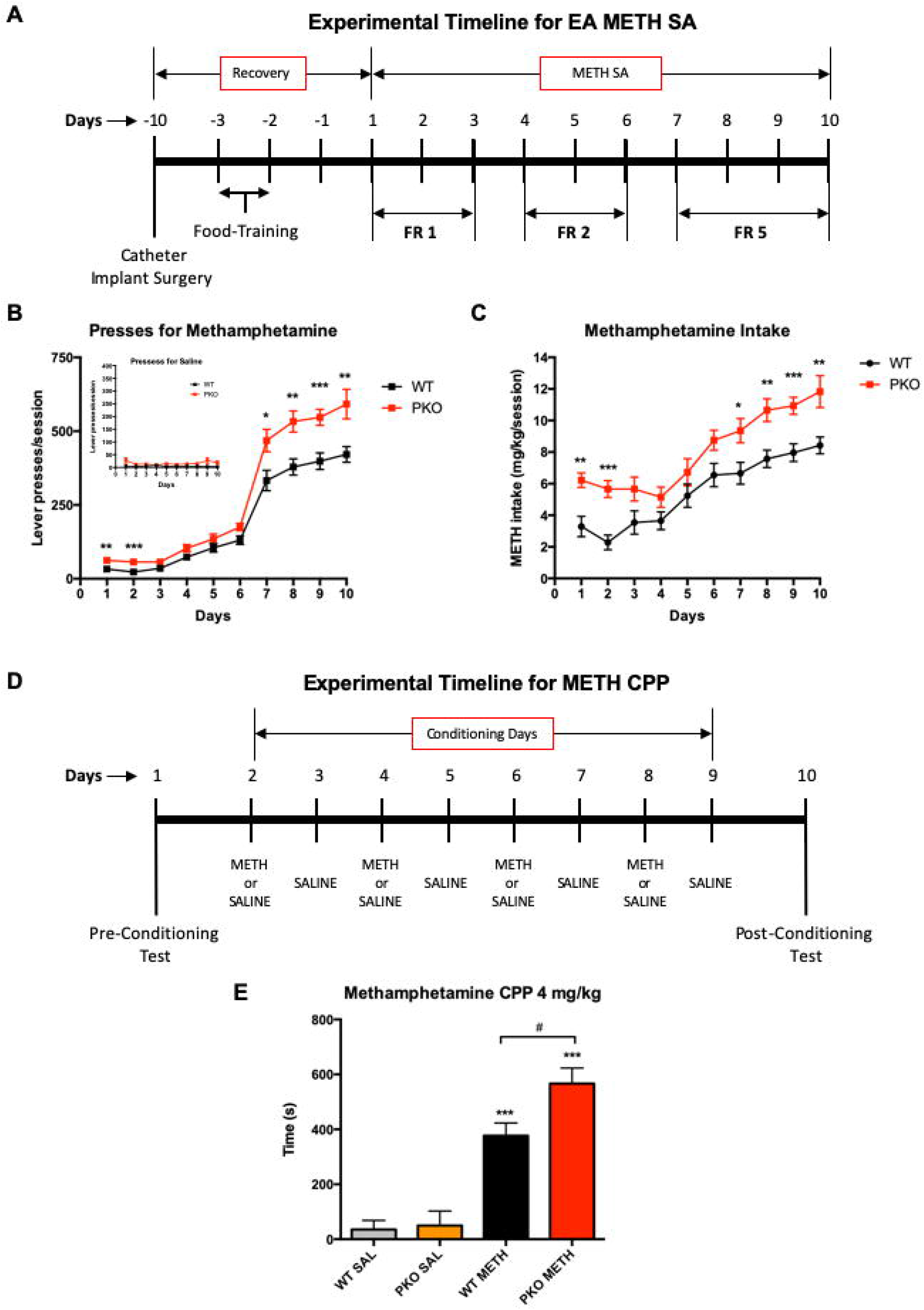
Parkin Knockout Increases Extended-Access Methamphetamine Self-Administration (EA METH SA) and METH Reward. (A) Timeline of EA METH SA experiment. (B, C) Escalation of METH taking in *parkin* gene knockout (PKO) rats as compared to wild-type (WT) rats during a progressive ratio schedule of reinforcement: FR1, FR2, and FR5. The PKO rats pressed for METH at significantly higher rate than the WT rats (B) with a consequent higher consumption of METH (C). There was no significant difference between PKO and WT rats in respect of number of presses for saline (B insert). \**p*<0.05, \*\**p*<0.01, \*\*\**p*<0.001, n=11/group. (D) Timeline of conditioned place preference experiment. (E) The PKO rats displayed a significantly higher preference for METH (4mg/kg) than the WT rats. \*\*\**p*<0.001 as compared to the respective saline (SAL) controls, ^#^*p*<0.05 WT *vs.* PKO, n=6/group. Data is expressed as mean ± SEM. Abbreviations: CPP, conditioned place preference; FR, fixed ratio.

**Figure 2.**
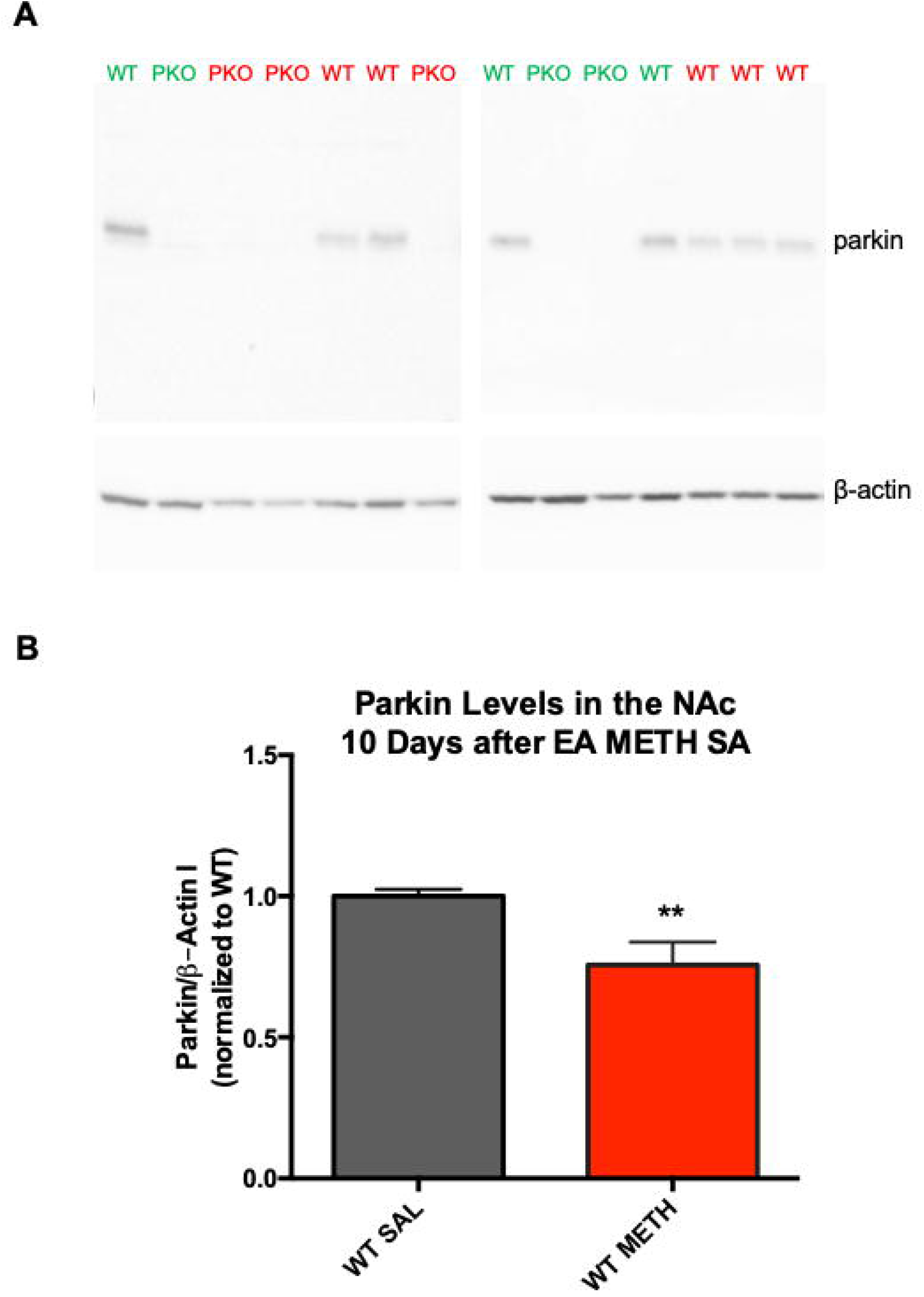
Extended-Access Methamphetamine Self-Administration (EA METH SA) Decreases Parkin Protein Levels in the Nucleus Accumbens. (A) Representative PARKIN bands (~52kDa) assessed in nucleus accumbens of wild type (WT) rats and *parkin* knockout (PKO) rats (to confirm loss of PARKIN protein), with β-ACTIN serving as a loading control. (B) Quantification of PARKIN bands. EA METH SA led to a deficit (−24%) in PARKIN levels in the nucleus accumbens of WT rats at 10d after the last operant session. \*\**p*<0.01, n=6/group. Data is expressed as mean ± SEM. Abbreviations: NAc, nucleus accumbens.

Supplementary Online Data provides extensive details on stereotaxic surgery as well as the EA METH SA and conditioned place preference.

### Electrophoresis and Western Blotting

Rats were sacrificed 10d after the last operant session. Nucleus accumbens was punched out of 2mm-thick coronal brain section encompassing 0.7-2.7mm from bregma. Electrophoresis and western blotting on nucleus accumbens pieces were performed as previously described^34^, utilizing anti-PARKIN and anti-β-ACTIN (1:1,000, 2132S and 3700S, respectively, Cell Signaling Technology, Danvers, MA, USA) primary antibodies. Immunoreactivities were quantified using ImageJ software (National Institutes of Health, Bethesda, MD, USA) and presented as relative optical densities normalized to control samples.

### Statistical Analyses

The experimental data sets were analyzed using IBM SPSS v.25 (IBM, Armonk, NY). EA METH SA data was analyzed with two-way mixed-design ANOVA whereas METH conditioned place preference data was analyzed with two-way factorial ANOVA, both with Bonferroni correction. Before each ANOVA, the data was examined for the presence of outliers using an IQR of 3. After the removal of outlier(s) the data was examined for normality with Shapiro-Wilk’s test, equality of variances (Levene’s test), and sphericity (Mauchly’s test). When Mauchly’s test was statistically significant, a Greenhouse-Geisser correction was employed. Group differences in PARKIN levels were assessed with the Student’s *t*-test. To assess sizes of genotype-mediated effects and probabilities of the presence of the effects, partial eta square (η_p_^2^) and power of analysis (1-β) values were calculated. Statistical significance was set at *p*≥0.05. All data, with the exception of body weight data, is reported as mean ± SEM. Body weight data is reported as mean ± SD (Suppl. Fig.S1).

## RESULTS

### Parkin Knockout Increases Extended-Access METH Self-Administration

To determine whether PKO rats are more dependent on METH as compared to WT rats, the rats from both genotypes were subjected to the EA METH SA test (Fig.1A). Both WT and PKO rats showed an escalation of METH self-administration (Fig.1B, C) The PKO rats weighed more than WT rats but lost weight at the same rate as WT rats during EA METH SA (Fig.S1A). Including body weight as a covariate in ANOVA analysis showed that it did not influence METH self-administration (*F*_(1,19)_=0.100, *p*=0.756, η_p_^2^=0.005, 1-β=0.06, n=11/genotype). There was a significant main effect of genotype (*F*_(1,20)_=16.7, *p*<0.01, η_p_^2^=0.454, 1-β=0.973) as well as time (*F*(2.9,58)=322, *p*<0.001, η_p_^2^=0.942, 1-β=1.0) on lever presses for METH, and significant interaction between the two factors (*F*_(2.9,58)_=5.4, *p*<0.01, η_p_^2^=0.213, 1-β=0.913). Pairwise comparisons revealed that PKOs pressed more than WTs during the first two sessions (*p*<0.01 and *p*<0.001, respectively) and during the compulsive phase of METH self-administration (day 7, *p*<0.05 and days 8-10, *p*<0.01) (Fig.1B). Concordant with this result, genotype and time both significantly influenced METH intake (genotype: *F*(1,20)=13.8. *p*<0.01, η_p_^2^=0.409, 1-β=0.941; time: *F*_(3.6,71)_=53.8, *p*<0.001, η_p_^2^=0.729, 1-β=1.0), but there was no significant interaction between the two factors (*F*_(3.6,71)_=1.40, *p*=0.246). The pairwise comparisons revealed that PKO rats self-administered significantly more METH during the first two sessions (*p*<0.01 and *p*<0.001, respectively) and during the compulsive phase of METH self-administration (day 7, *p*<0.05 and days 8-10, *p*<0.01) (Fig.1C). There was no significant difference between PKO and WT rats in respect to number of presses for saline (Fig.1B insert, n=4/genotype), confirming that METH, not the light stimulus, acted as a reinforcer.

The results from the EA METH SA indicate that METH exerts stronger reinforcing effects in rats lacking PARKIN than in WT controls. The effect of PARKIN knockout on METH self-administration was of large size (large η_p_^2^) and significant, thus strongly suggesting PARKIN as a suppressor of reinforcing effects of METH.

### Parkin Knockout Increases Preference for METH-Paired Compartment

Compulsive drug use is maintained by the rewarding properties of the drug and by the loss of control over drug intake. To determine whether PARKIN knockout increases the rewarding effects of METH, PKO and WT rats underwent the conditioned place preference test (Fig.1D). Though the conditioned place preference test has limitations^35^, it provides information about the rewarding effect of contextual cues associated with a drug stimulus. Two-way ANOVA revealed a trend for significant interaction between genotype and treatment (*F*_(1,20)_=3.31, *p*=0.084, η_p_^2^=0.142, 1-β=0.410), and main effects of both genotype and treatment on METH preference, with the effect of treatment being stronger than the effect of genotype (*F*_(1,20)_=79.2, *p*<0.001, η_p_^2^=0.798, 1-β=1.00 and *F*_(1,20)_=4.50, *p*<0.05, η_p_^2^=0.184, 1-β=0.523, respectively, n=6/group). Pairwise comparisons revealed that there was no significant difference in baseline preference for a compartment between PKO and WT rats (*p*=0.834) (Fig.1E). Both genotypes showed a significant METH-induced conditioned place preference (*p*<0.001), with the PKO rats displaying significantly higher preference for METH-paired compartment than the WT rats (*p*<0.05) (Fig.1F). ANCOVA determined that body weight had no influence on the results (not shown).

The results from the conditioned place preference test suggest that METH-paired contextual cues are more rewarding for PKO rats than for WT rats.

### Extended-Access METH Self-Administration Leads to Parkin Deficit in the Nucleus Accumbens

We previously determined that METH binge decreases PARKIN levels in the striatum via oxidative damage^36^. To determine whether exposure of WT rats to EA METH SA paradigm decreases PARKIN levels in the nucleus accumbens, PARKIN levels were assessed in rats that self-administered METH at 10 days after the last operant session and compared to PARKIN levels in rats exposed to saline. The nucleus accumbens tissue samples from PKO rats were included in the analysis to ascertain the lack of PARKIN protein in these samples. METH-exposed rats had significantly lower levels of accumbal PARKIN compared to saline-exposed rats (−24%, *t*=3.02, *df*=15, *p*<0.01, n=8-9) (Fig.2B).

### Validation of Parkin Overexpression

Viral vector microinjections took place 3 weeks before METH self-administration or conditioned place preference experiments to allow for maximal PARKIN overexpression (Fig.3A). Investigation of the microinjection sites using Trypan blue dye confirmed that during microinjections, the needle reached approximately the middle of the nucleus accumbens (Fig.3B) and that 2μL of the dye did not spread beyond the borders of the nucleus accumbens (Fig.3B, C). Administration of 2×10^7^ or 6.5×10^7^ TUs/side AAV2/6-parkin consistently produced a 4-6-fold and 15-20-fold increase in PARKIN levels, respectively, in WT rats (Fig.3D). Parkin overexpression lasted till the end of the behavioral experiments (~4 weeks) (Fig.3E).

**Figure 3.**
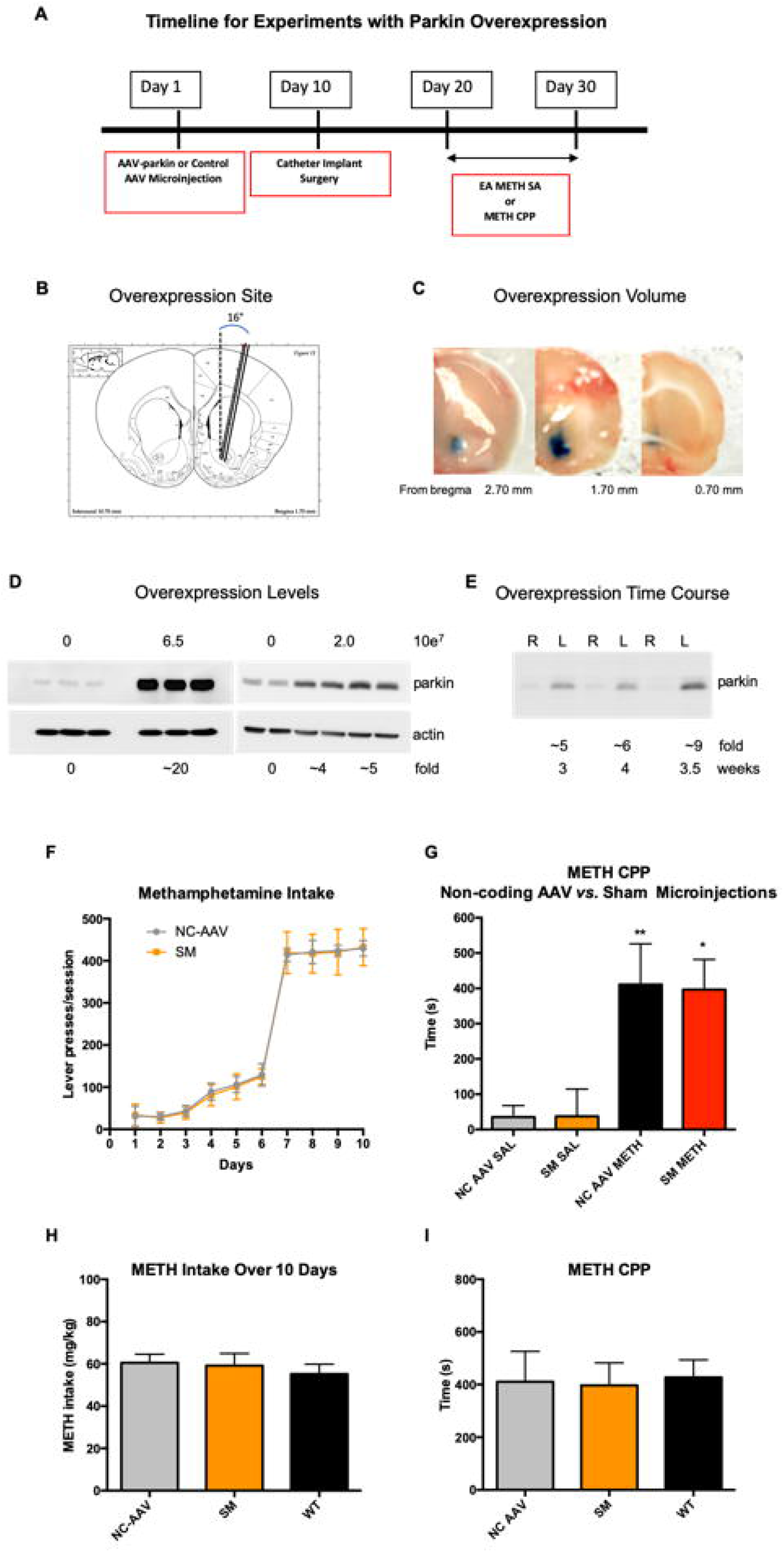
Validation of Parkin Overexpression. (A) Timeline for experiments with PARKIN overexpression. (B, C) Validation of the overexpression site (nucleus accumbens, 0.7-2.7mm from bregma) and spread (medial-lateral, dorsal-ventral and anterior-posterior). (D) Two levels of PARKIN overexpression 2×10^7^ or 6.5×10^7^ TUs/side in the nucleus accumbens (n=3-4/group). (E) Time course of PARKIN overexpression (2×10^7^) showing that excess of PARKIN persists at least for duration of the study (~4 weeks). (F-I) Neither non-coding AAV2/6 (NC-AAV) nor sham microinjection (SM) had an effect on methamphetamine (METH) self-administration or preference for METH-paired compartment in the conditions place preference test in wild type (WT) rats. (F) Lever presses in NC-AAV and SM group. (G) Conditioned place preference test results for NC-AAV and SM group. (H) METH intake in WT *vs.* NC-AAV and SM rats. (I) Preference for METH-paired compartment in WT *vs.* NC-AAV and SM rats. Data is expressed as mean ± SEM. Abbreviations: CPP, conditioned place preference; NAc, nucleus accumbens; L, left; R, right.

We next assessed whether non-coding AAV2/6 or microinjection surgery has an effect on the outcome of EA METH SA or METH conditioned place preference test. Rats were microinjected with non-coding AAV or underwent sham microinjections with 2uL saline. These groups were named Non-Coding AAV (NC-AAV) group and Sham-Microinjection (SM) group. Both groups underwent EA METH SA and METH conditioned place preference test. During METH self-administration, both NC-AAV and SM rats pressed for METH at a similar rate (main effect of time: *F*_(9,54)_=62.2, *p*=0.06, η_p_^2^=0.912, 1-β=1.0, n=4/group; main effect of surgery type: *F*_(1, 12)_=, *p*=0.862, η_p_^2^=0.005, 1-β=0.053) (Fig.3F). During the conditioned place preference test, both NC-AAV and SM rats displayed a preference for the METH-paired compartment (main effect of treatment: *F*_(1,12)_=19.6, *p*<0.001, η_p_^2^= 0.620, 1-β=0.982, n=4/group; NC-AAV: *F*_(1,12)_=10.2, *p*<0.01, η_p_^2^=0.460, 1-β=0.835; SM: *F*_(1,12)_=9.36, *p*<0.05, η_p_^2^= 0.438. 1-β=0.802) and did not differ in respect to time spent in this compartment (*p*>0.1) (Fig.3G). Similarly, these experimental groups did not significantly differ in respect to time spent in the saline-paired compartment (*p*>0.1). There was no statistically significant interaction between treatment and different microinjections on METH-paired place preference (*p*>0.1). The NC-AAV and SM groups did not differ from non-microinjected WT rats in respect to self-administered METH or degree of preference for the METH-paired compartment. Comparison of the three conditions are summarized in Fig.3H and Fig.3I. This data indicated that neither non-coding AAV2/6 nor sham microinjections influenced the outcomes of EA METH SA or METH conditioned place preference test.

### Wild-type Rats Overexpressing PARKIN Self-administer Less METH and Spend Less Time in METH-Paired Compartment than Wild-type Controls

Since WT rats subjected to EA METH SA had lower PARKIN levels in the nucleus accumbens than saline controls, we next investigated whether overexpression of PARKIN in the nucleus accumbens of WT rats would result in attenuated METH self-administration as compared to non-overexpressing WT rats. Two concentrations of PARKIN-AAV2/6 were utilized to overexpress PARKIN - 2×10^7^ or 6.5×10^7^ TUs/side (4-6 or 15-20-fold excess of PARKIN). The PO rats weighed slightly less than WT controls (Fig.S1B); however, body weight did not have a significant effect on METH self-administration as a covariate (*F*_(1,9)_=0.006, *p*=0.939, η_p_^2^=0.001, 1-β=0.051, n=6/group). At low levels of PARKIN overexpression, there was a significant main effect of time (*F*_(3.1,31.5)_=51.6, *p*<0.001, η_p_^2^=0.838, 1-β=1.0), but not of genotype (*p*>0.1), on lever presses for METH (Fig.4A). There was no statistically significant interaction between the two factors (*p*>0.1). Statistical analysis of METH intake over 10 days produced similar a result: there was a significant effect of time (*F*_(1.54,15.4)_=187, *p*<0.001, η_p_^2^=0.949, 1-β=1.0), but no significant main effect of genotype (*p*>0.1) and no significant genotype x time interaction (*p*>0.1) (Fig.4B). At high levels of PARKIN overexpression, there was a strong significant effect of genotype *F*(1,10)=47.8, *p*<0.0001, η_p_^2^=0.827, 1-β=1.0), as well as time (*F*_(2.19,21.9)_=543, *p*<0.0001, η_p_^2^=0.982, 1-β=1.0), on METH lever pressing (Fig.4C) and significant interaction between these variables (*F*_(2.19,21.9)_=34.9, *p*<0.0001, η_p_^2^=0.577). Pairwise comparisons revealed that PKO rats pressed significantly less for METH on days 4-10, with the strongest effect at the compulsive stage of METH abuse (day 4, *p*<0.001; days 5 and 6, *p*<0.05; day 7 *p*<0.001; days 8-10, *p*<0.0001) (Fig.4C). In concordance, there was a strong significant effect of genotype (*F*_(1,10)_=31.6, *p*<0.0001, η_p_^2^=0.760, 1-β=0.999) as well as time (*F*_(4,40)_=103, *p*<0.0001, η_p_^2^=0.912, 1-β=1.0) on METH intake and significant interaction between these variables (*F*_(4,40)_=14.4, *p*<0.0001, η_p_^2^=0.573). As with lever presses, pairwise comparisons showed that PKO took significantly less METH on days 4-10, with the strongest effect on days 8-10 (day 4, *p*<0.001; days 5 and 6, *p*<0.05; day 7, *p*<0.001; days 8-10, *p*<0.0001) (Fig.4D). Figure 4E presents a comparison of METH intake during the compulsive phase of EA METH SA (days 8-10).

**Figure 4.**
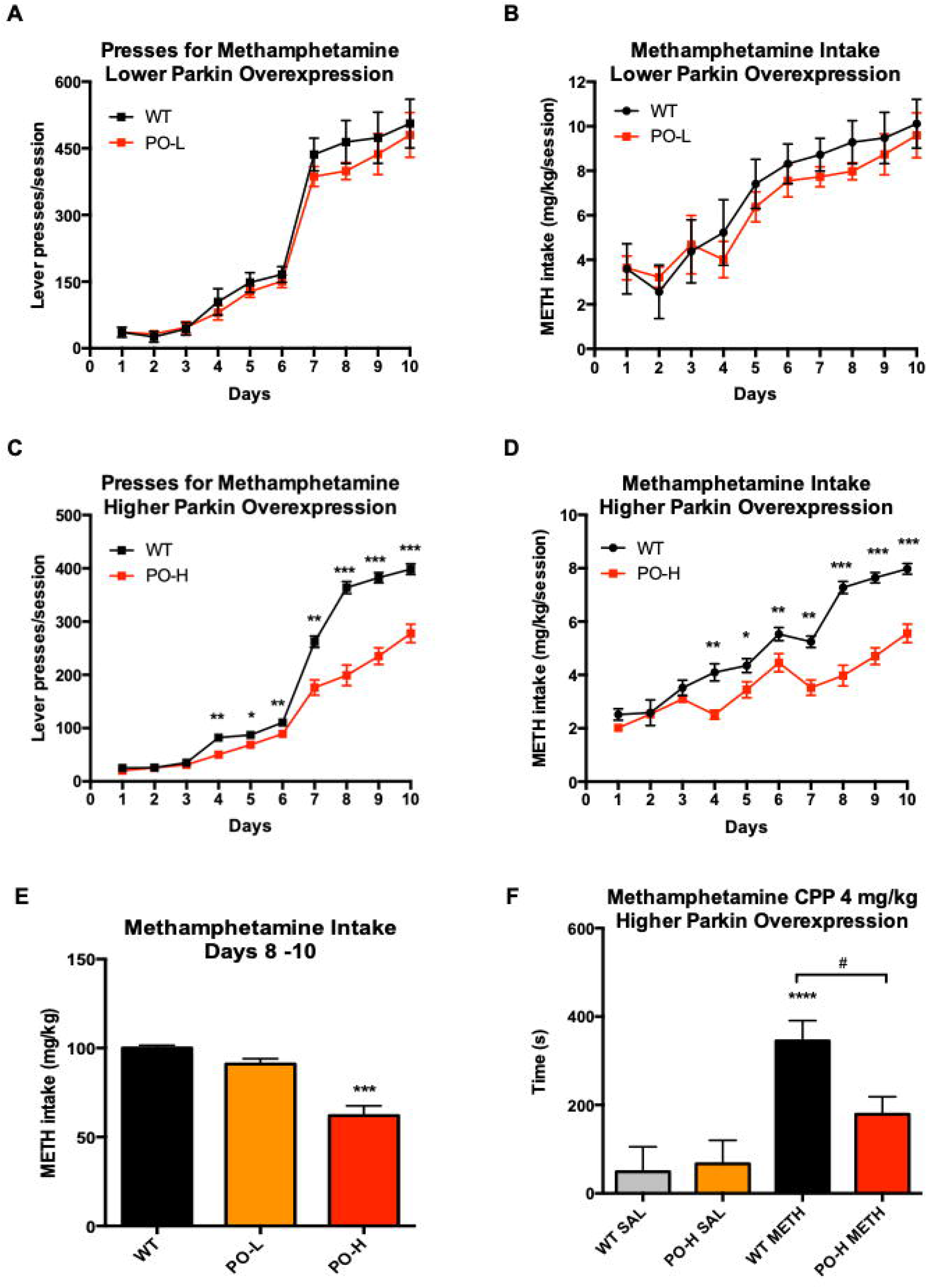
Parkin Overexpression in the Nucleus Accumbens of WT Rats Attenuates Extended-Access Methamphetamine Self-Administration (EA METH SA) and METH Reward. Escalation of METH taking in rats overexpressing PARKIN in the nucleus accumbens as compared to non-expressing wild-type (WT) rats during a progressive ratio schedule of reinforcement: FR1, FR2, and FR5. (A) At the low PARKIN overexpression (4-6-fold), the parkin overexpressing (PO) rats did not press for METH at a significantly lower rate than the WT rats and, consequently, (B) they did not consume more METH than the WT rats. (C) At the high parkin overexpression (15-20-fold), the PO rats did press for METH at a significantly lower rate than the WT rats and (D) consumed less METH than the WT rats. (E) Quantification of compulsive METH intake during days 8-10. \**p*<0.05, \*\**p*<0.01, \*\*\**p*<0.001, WT *vs.* PO-H, n=6/group. (F) The 10-15-fold excess of PARKIN was sufficient to significantly attenuate preference for METH (4mg/kg) in WT rats in the conditioned place preference test. ***p<0.001 as compared to the respective saline controls, ^#^*p*<0.05 WT *vs.* PO-H, n=6/group. Data is expressed as mean ± SEM. Abbreviations: NAc, nucleus accumbens; FR, fixed ratio; PO-L, lower PAKIN overexpression; PO-H, higher PARKIN overexpression; SAL, saline.

WT rats overexpressing PARKIN at high levels spent less time in METH-paired compartment in the conditioned preference test than non-overexpressing WT controls (Fig.4F). Two-way ANOVA revealed the main effect of treatment on METH preference (*F*_(1,20)_=17.1, *p*<0.001, η_p_^2^=0.461, 1-β=0.976) and a trend for significant interaction between genotype and treatment (*F*_(1,20)_=3.47, *p*=0.077, η_p_^2^=0.148, 1-β=0.426). Pairwise comparisons detected no significant difference in baseline preference for a compartment between PO and WT rats (*p*=0.800, n=6/group). Significant preference for the METH-paired compartment, as compared to the saline-paired compartment, was detected in WT rats (*p*<0.0001) but not in PO rats (*p*=0.123), however, there was a difference between genotypes in METH group (*p*<0.05) (Fig.4F). The ANCOVA analysis determined that body weight had no influence on the results (not shown).

These results further support the conclusion that PARKIN negatively regulates reinforcing/rewarding properties of METH.

### Parkin Knockout Rats Overexpressing PARKIN Self-administer Less METH During the First Half of Drug Self-administration Compared to PARKIN-deficient Rats

The next experiment aimed to determine whether overexpression of PARKIN in the nucleus accumbens of PKO rats would attenuate METH self-administration as it did in nucleus accumbens of WT rats. High levels (15-20-fold) of PARKIN were overexpressed in the nucleus accumbens of PKO rats (n=6/group), while control rats were microinjected with saline (sham microinjection surgery). PKO and PKO PO rats did not differ in respect to body weight during the 10 days of METH SA (Supp. Fig.1C). There was a significant main effect of time (*F*_(1.2,11)_=192, *p*<0.0001, η_p_^2^=0.838, 1-β=1.0) and a weaker main effect of genotype (*F*_(1,9)_=5.15, *p*=0.05, η_p_^2^=0.364, 1-β=0.525) on lever presses for METH between the genotypes (Fig.5A). There was no statistically significant interaction between the time and genotype (*p*>0.1). Examination of METH intake data produced similar results: a significant main effect of time (*F*(1.5,13.4)=29.3, *p*<0.0001 0.765, 1-β=1.0) and genotype (*F*_(1,9)_=18.3, *p*<0.01, η_p_^2^=0.671, 1-β=0.966) but no significant genotype x time interaction (*p*>0.1). Pairwise comparisons revealed that PKO PO rats pressed significantly less for METH and, consequently, ingested significantly less of the drug than non-overexpressing PKO rats in the first half of EA METH SA (days 1, 4 and 5) (*p*<0.01, p<0.01, and *p*<0.001, respectively) but not at later stages of EA METH SA (Fig.5B).

**Figure 5.**
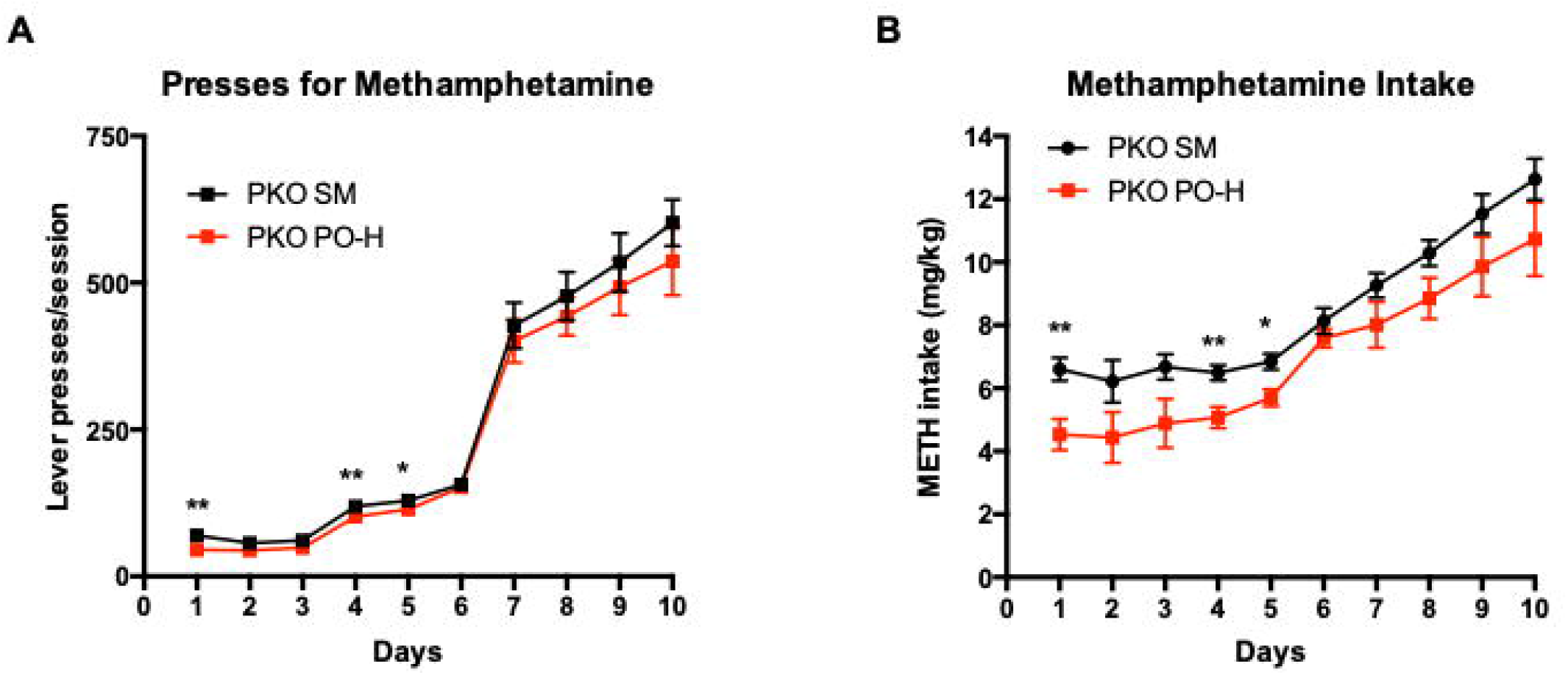
The Effect of Parkin Overexpression in the Nucleus Accumbens of *Parkin* Gene Knockout (PKO) Rats. High PARKIN overexpression (15-20-fold) in the nucleus accumbens of *parkin* knockout PKO rats significantly attenuated methamphetamine (METH) taking in the first half of the extended-access METH self-administration as compared to non-overexpressing PKO rats. (A) Lever presses for METH, (B) METH intake. \**p*<0.05, \*\**p*<0.01, n=5-6/group. Data is expressed as mean ± SEM.

This result suggests that PARKIN knockout-induced compensatory changes may have lessen the effect of PARKIN overexpression.

## DISCUSSION

METH Use Disorder is a disorder without a pharmacotherapy and, therefore, in need of new drug targets for development of new drugs. In this study, we used genetically engineered rats to evaluate PARKIN as a potential target for METH Use Disorder. We found that PARKIN knockout rats self-administered more METH and spent more time in the METH-paired environment than WT rats whereas WT rats overexpressing PARKIN in the nucleus accumbens self-administered less METH and spent less time in the METH-paired environment. We also found that PARKIN knockout rats overexpressing PARKIN in the nucleus accumbens self-administered less METH during the first half of drug self-administration days than PARKIN-deficient rats. The results indicate that rats with PARKIN excess or PARKIN deficit are useful models for studying neural substrates underlying “resilience” or vulnerability to METH Use Disorder, and identify PARKIN as a novel potential drug target to treat heavy use of METH.

### The Role of PARKIN in METH-Taking Behavior

Drug-taking behavior is a consequence of reinforcing effects of the drug of abuse, mediated by goal-directed and habit systems in the brain. At the initial stages of drug self-administration, the goal-directed system directs the relationship between action (lever pressing) and outcome (drug infusion and experiencing sought-after drug effects). After prolonged drug taking, instrumental action becomes compulsive and habitual^37^. Escalation of drug self-administration is a hallmark of compulsive drug taking and drug dependence^32^. Hence, escalation of METH self-administration by both PKO and WT rats despite increasing effort required to obtain METH (from 1 to 5 presses needed to obtain METH injection) suggests increasing motivation to obtain the drug, leading to METH dependence. Compared to the WT controls, the PKO rats pressed significantly more for METH and, consequently, consumed more METH than WT rats during the initial stage (FR1) and during the FR5 stage (compulsive intake) of EA METH SA. In contrast, the PO rats pressed significantly less for METH and consumed more of the drug at these stages. Two conclusions can be reached based on these findings. First is that PARKIN may be involved in negative modulation of reinforcing/rewarding effects of METH within the nucleus accumbens, which is the main brain region mediating rewarding/reinforcing effects^38^. This conclusion is supported by the results from the conditioned place preference test. The conditioned place preference test and self-administration measure different processes, namely drug reward and context-drug associative learning *vs.* acquisition of a drug reinforced operant response, respectively. However, nucleus accumbens involvement and reward are common and there is reasonable concordance between drugs that produce conditioned place preference and drugs that are self-administered^35^. Because both paradigms involve associative learning^35,39^ and because the nucleus accumbens plays a role in this type of learning^37,40,41^, PARKIN may have a role in processes associating METH with stimuli and context. In addition to the aforementioned functions, the nucleus accumbens controls goal-directed behavior during instrumental conditioning^37,42–45^. Consequently, the second conclusion is that accumbal PARKIN may negatively regulate goal-directed behavior during METH self-administration. We demonstrated that the volume spread after the microinjections was largely confined to the nucleus accumbens. Therefore, it is unlikely that the difference in METH self-administration observed between PO and WT rats was due to unspecific PARKIN overexpression in the adjacent dorsomedial striatum, which is a key brain area regulating goal-directed behavior^37,46–48^, but rather to overexpression of PARKIN in the nucleus accumbens, which has a role in action selection^45^. Habitual drug taking is mediated by the dorsolateral striatum^37,46,49,50^ and it is not known whether striatal PARKIN affects this behavior. Overexpression of PARKIN in the nucleus accumbens likely resulted in its excess not only in accumbal perikarya, but also in terminals located in brain areas to which the nucleus accumbens projects to^33^. Consequently, PARKIN could have presynaptically modulated METH taking in these brain areas. Even though PKO and PO rats displayed opposite addictive behaviors, molecular mechanisms underlying these behaviors may be different due to compensatory mechanisms that occurred in the brains of PKO but not in PO rats during development^51^. Such changes might explain the smaller effect of PARKIN overexpression on METH taking in PKO PO rats as compared to PO rats.

### Molecular Determinants of Altered METH-Taking Behavior in PKO and PO Rats

In relation to the present investigation, studies in young adult PKO mice (3–4-month-old) reported no change in the levels of striatal dopamine or dopamine receptors as compared to WT littermates^52,53^, with dopamine metabolites reported unchanged or elevated^54,55^. Our previous study in young adult PKO rats (2-month-old) similarly found no change in dopamine and dopamine metabolites but reported that PKO rats had lower levels of glycosylated form of postsynaptic D2 receptor than WT rats in the dorsal striatum^34^. Low D2/D3 receptor availability in striatal regions, including the nucleus accumbens, is considered a risk factor for METH Use Disorder because it is a molecular feature of impulsivity; animal as well as human data support this notion^56,57^. Furthermore, it has been demonstrated that D2-expressing medium spiny neurons in the nucleus accumbens antagonize the reinforcing and rewarding effects of another psychostimulant, cocaine^58^. *Parkin* gene knockout results in augmented glutamatergic neurotransmission and altered levels of glutamate receptors as well as GABAB receptors in mice^31^. This data suggests the involvement of PARKIN protein in regulation of these receptors. In conclusion, dopamine, glutamate and/or γ-aminobutyric acid (GABA) likely modulate the rewarding/reinforcing effects of METH *via* their receptors^59–64^.

METH Use Disorder has been linked to alterations in energy metabolism and cytoskeletal arrangement as well as to oxidative stress and inflammation^11–21,65–67^. For example, METH-induced inflammation in the nucleus accumbens was shown to mediate METH reinforcing properties^11,65^. In the EA METH SA paradigm, large doses of METH were self-administered, likely causing oxidative stress and inflammation in multiple brain areas, including the nucleus accumbens^11,68^. PARKIN has been linked to energy metabolism, and cytoskeletal arrangement as well as to protection against oxidative stress, and inflammation^25,26,29–31^. Therefore, PARKIN is well-positioned to regulate responses to METH and could have decreased METH intake in rats via any of the aforementioned mechanisms.

Addiction circuitry includes multiple brain areas^69^, all of which have likely undergone neuroadaptive changes in the PKO rats during development^51^. We hypothesize that because of these changes and lack of PARKIN in addiction circuitry, overexpression of PARKIN in the nucleus accumbens only had smaller effect on METH self-administration in PKO rats than WT rats. Overexpression of PARKIN in the nucleus accumbens and prefrontal cortex and/or dorsal striatum may be necessary to suppress the addictive phenotype in PKO rats. Data from PKO PO rats suggest that PARKIN has stronger effect on molecular mechanisms underlying reinforcing properties than on those underlying loss of control over METH intake in these rats. It remains to be determined whether PARKIN overexpression in the nucleus accumbens of PKO rats would have more pronounced effect on METH self-administration in a paradigm with short-access to the drug (e.g. 1h), consequently leading to low METH intake.

### Limitations and Methodological Considerations

A limitation of using PKO rats is that developmental adaptations to the absence of the *parkin* gene may have compensated for an early loss of gene function. As a result, such adaptations may have obscured the analysis of the effects of PARKIN protein *per se*. Nevertheless, overexpression of PARKIN resulted in behaviors opposite to those observed in PKO rats, thus validating the role of PARKIN in METH self-administration. High level of PARKIN overexpression was needed for a desired effect, which can be viewed as non-translational. However, since we employed a model of very high METH intake, lower levels of PARKIN overexpression will likely work with models of moderate METH intake.

### Conclusions

To date, pharmacological approaches have failed as treatments for METH Use Disorder. We demonstrated for the first time that PARKIN plays important role in this disorder. Specifically, we demonstrated that PARKIN modulates the rewarding/reinforcing properties of high-dose METH. The role of PARKIN in drug use disorder has not been previously studied; therefore, our study is novel. From the public health point of view, the most important finding of this investigation is that overexpression of PARKIN in the nucleus accumbens alone was sufficient to attenuate intake of very high doses of METH by WT rats, making PARKIN a novel potential drug target to treat heavy use of METH. In addition, we demonstrated that rats with PARKIN excess or PARKIN deficit are useful models for studying neural substrates underlying “resilience” or vulnerability to METH Use Disorder.

## Acknowledgements

This research was funded by Wayne State University and partially by NIH/NIDA DA034738 grant.

## Author Contributions

A.S. and A.M. analyzed and interpreted the data, prepared the figures, and co-wrote the manuscript. A.H. significantly contributed to data generation. B.S. provided AAVs and consulted on PARKIN overexpression. All authors reviewed and edited the manuscript.

## REFERENCES

1 Jalal, H. et al. Changing dynamics of the drug overdose epidemic in the United States from 1979 through 2016. Science 361, (2018).

2 NIDA. Overdose Death Rates. (2018).

3 Rawson, R. A. Current research on the epidemiology, medical and psychiatric effects, and treatment of methamphetamine use. J Food Drug Anal 21, S77–S81, (2013).

4 Courtney, K. E. & Ray, L. A. Methamphetamine: an update on epidemiology, pharmacology, clinical phenomenology, and treatment literature. Drug Alcohol Depend 143, 11–21, (2014).

5 Karila, L. et al. Pharmacological approaches to methamphetamine dependence: a focused review. Br J Clin Pharmacol 69, 578–592, (2010).

6 Logan, B. K. Methamphetamine - Effects on Human Performance and Behavior. Forensic Sci Rev 14, 133–151, (2002).

7 Stock, A. K., Radle, M. & Beste, C. Methamphetamine-associated difficulties in cognitive control allocation may normalize after prolonged abstinence. Prog Neuropsychopharmacol Biol Psychiatry 88, 41–52, (2019).

8 Rusyniak, D. E. Neurologic manifestations of chronic methamphetamine abuse. Neurol Clin 29, 641–655, (2011).

9 Brecht, M. L., Lovinger, K., Herbeck, D. M. & Urada, D. Patterns of treatment utilization and methamphetamine use during first 10 years after methamphetamine initiation. J Subst Abuse Treat 44, 548–556, (2013).

10 UNODC. (Vienna, 2018).

11 Jang, E. Y. et al. The role of reactive oxygen species in methamphetamine self-administration and dopamine release in the nucleus accumbens. Addict Biol 22, 1304–1315, (2017).

12 Najera, J. A. et al. Methamphetamine abuse affects gene expression in brain-derived microglia of SIV-infected macaques to enhance inflammation and promote virus targets. BMC Immunol 17, 7, (2016).

13 Yamamoto, B. K. & Raudensky, J. The role of oxidative stress, metabolic compromise, and inflammation in neuronal injury produced by amphetamine-related drugs of abuse. J Neuroimmune Pharmacol 3, 203–217, (2008).

14 Stephans, S. E., Whittingham, T. S., Douglas, A. J., Lust, W. D. & Yamamoto, B. K. Substrates of energy metabolism attenuate methamphetamine-induced neurotoxicity in striatum. J Neurochem 71, 613–621, (1998).

15 Wang, G. J. et al. Partial recovery of brain metabolism in methamphetamine abusers after protracted abstinence. Am J Psychiatry 161, 242–248, (2004).

16 Volkow, N. D. et al. Recovery of dopamine transporters with methamphetamine detoxification is not linked to changes in dopamine release. Neuroimage 121, 20–28, (2015).

17 Sekine, Y. et al. Methamphetamine causes microglial activation in the brains of human abusers. J Neurosci 28, 5756–5761, (2008).

18 Young, E. J., Briggs, S. B. & Miller, C. A. The Actin Cytoskeleton as a Therapeutic Target for the Prevention of Relapse to Methamphetamine Use. CNS Neurol Disord Drug Targets 14, 731–737, (2015).

19 Moussawi, K. & Kalivas, P. W. Group II metabotropic glutamate receptors (mGlu2/3) in drug addiction. Eur J Pharmacol 639, 115–122, (2010).

20 Jayanthi, S. et al. Methamphetamine downregulates striatal glutamate receptors via diverse epigenetic mechanisms. Biol Psychiatry 76, 47–56, (2014).

21 Krasnova, I. N., Justinova, Z. & Cadet, J. L. Methamphetamine addiction: involvement of CREB and neuroinflammatory signaling pathways. Psychopharmacology (Berl) 233, 1945–1962, (2016).

22 Ernst, T., Chang, L., Leonido-Yee, M. & Speck, O. Evidence for long-term neurotoxicity associated with methamphetamine abuse: A 1H MRS study. Neurology 54, 1344–1349, (2000).

23 Chan, P. et al. Rapid ATP loss caused by methamphetamine in the mouse striatum: relationship between energy impairment and dopaminergic neurotoxicity. J Neurochem 62, 2484–2487, (1994).

24 Ruan, Q. T. et al. A Mutation in Hnrnph1 That Decreases Methamphetamine-Induced Reinforcement, Reward, and Dopamine Release and Increases Synaptosomal hnRNP H and Mitochondrial Proteins. J Neurosci 40, 107–130, (2020).

25 Sassone, J. et al. The synaptic function of parkin. Brain 140, 2265–2272, (2017).

26 Seirafi, M., Kozlov, G. & Gehring, K. Parkin structure and function. FEBS J 282, 2076–2088, (2015).

27 Jiang, H. et al. Parkin controls dopamine utilization in human midbrain dopaminergic neurons derived from induced pluripotent stem cells. Nat Commun 3, 668, (2012).

28 Moszczynska, A. et al. Parkin disrupts the alpha-synuclein/dopamine transporter interaction: consequences toward dopamine-induced toxicity. J Mol Neurosci 32, 217–227, (2007).

29 Huynh, D. P., Scoles, D. R., Ho, T. H., Del Bigio, M. R. & Pulst, S. M. Parkin is associated with actin filaments in neuronal and nonneural cells. Ann Neurol 48, 737–744, (2000).

30 Ren, Y., Zhao, J. & Feng, J. Parkin binds to alpha/beta tubulin and increases their ubiquitination and degradation. J Neurosci 23, 3316–3324, (2003).

31 Cremer, J. N. et al. Changes in the expression of neurotransmitter receptors in Parkin and DJ-1 knockout mice--A quantitative multireceptor study. Neuroscience 311, 539–551, (2015).

32 Edwards, S. & Koob, G. F. Escalation of drug self-administration as a hallmark of persistent addiction liability. Behav Pharmacol 24, 356–362, (2013).

33 Liu, B., Traini, R., Killinger, B., Schneider, B. & Moszczynska, A. Overexpression of parkin in the rat nigrostriatal dopamine system protects against methamphetamine neurotoxicity. Exp Neurol, (2013).

34 Gemechu, J. M. et al. Characterization of Dopaminergic System in the Striatum of Young Adult Park2(-/-) Knockout Rats. Sci Rep 8, 1517, (2018).

35 Bardo, M. T. & Bevins, R. A. Conditioned place preference: what does it add to our preclinical understanding of drug reward? Psychopharmacology (Berl) 153, 31–43, (2000).

36 Moszczynska, A. & Yamamoto, B. K. Methamphetamine oxidatively damages parkin and decreases the activity of 26S proteasome in vivo. J Neurochem 116, 1005–1017, (2011).

37 Everitt, B. J. & Robbins, T. W. Neural systems of reinforcement for drug addiction: from actions to habits to compulsion. Nat Neurosci 8, 1481–1489, (2005).

38 Scofield, M. D. et al. The Nucleus Accumbens: Mechanisms of Addiction across Drug Classes Reflect the Importance of Glutamate Homeostasis. Pharmacol Rev 68, 816–871, (2016).

39 Wise, R. A. Dopamine, learning and motivation. Nat Rev Neurosci 5, 483–494, (2004).

40 Di Chiara, G. Nucleus accumbens shell and core dopamine: differential role in behavior and addiction. Behav Brain Res 137, 75–114, (2002).

41 Whitaker, L. R. et al. Associative Learning Drives the Formation of Silent Synapses in Neuronal Ensembles of the Nucleus Accumbens. Biol Psychiatry 80, 246–256, (2016).

42 Mannella, F., Gurney, K. & Baldassarre, G. The nucleus accumbens as a nexus between values and goals in goal-directed behavior: a review and a new hypothesis. Front Behav Neurosci 7, 135, (2013).

43 Schultz, W., Apicella, P., Scarnati, E. & Ljungberg, T. Neuronal activity in monkey ventral striatum related to the expectation of reward. J Neurosci 12, 4595–4610, (1992).

44 Goto, Y. & Grace, A. A. Dopaminergic modulation of limbic and cortical drive of nucleus accumbens in goal-directed behavior. Nat Neurosci 8, 805–812, (2005).

45 Khamassi, M. & Humphries, M. D. Integrating cortico-limbic-basal ganglia architectures for learning model-based and model-free navigation strategies. Front Behav Neurosci 6, 79, (2012).

46 Yin, H. H. & Knowlton, B. J. The role of the basal ganglia in habit formation. Nat Rev Neurosci 7, 464–476, (2006).

47 Yin, H. H., Ostlund, S. B., Knowlton, B. J. & Balleine, B. W. The role of the dorsomedial striatum in instrumental conditioning. Eur J Neurosci 22, 513–523, (2005).

48 Yin, H. H., Knowlton, B. J. & Balleine, B. W. Blockade of NMDA receptors in the dorsomedial striatum prevents action-outcome learning in instrumental conditioning. Eur J Neurosci 22, 505–512, (2005).

49 Yin, H. H., Knowlton, B. J. & Balleine, B. W. Lesions of dorsolateral striatum preserve outcome expectancy but disrupt habit formation in instrumental learning. Eur J Neurosci 19, 181–189, (2004).

50 Yin, H. H., Knowlton, B. J. & Balleine, B. W. Inactivation of dorsolateral striatum enhances sensitivity to changes in the action-outcome contingency in instrumental conditioning. Behav Brain Res 166, 189–196, (2006).

51 Kreiner, G. Compensatory mechanisms in genetic models of neurodegeneration: are the mice better than humans? Front Cell Neurosci 9, 56, (2015).

52 Oyama, G. et al. Impaired in vivo dopamine release in parkin knockout mice. Brain Res 1352, 214–222, (2010).

53 Kitada, T. et al. Impaired dopamine release and synaptic plasticity in the striatum of parkin−/− mice. J Neurochem 110, 613–621, (2009).

54 Zhu, X. R. et al. Non-motor behavioural impairments in parkin-deficient mice. Eur J Neurosci 26, 1902–1911, (2007).

55 Menendez, J. et al. Suppression of Parkin enhances nigrostriatal and motor neuron lesion in mice over-expressing human-mutated tau protein. Hum Mol Genet 15, 2045–2058, (2006).

56 Lominac, K. D. et al. Mesocorticolimbic monoamine correlates of methamphetamine sensitization and motivation. Front Syst Neurosci 8, 70, (2014).

57 London, E. D. Impulsivity, Stimulant Abuse, and Dopamine Receptor Signaling. Adv Pharmacol 76, 67–84, (2016).

58 Baik, J. H. Dopamine signaling in reward-related behaviors. Front Neural Circuits 7, 152, (2013).

59 Arias-Carrion, O. et al. Orquestic regulation of neurotransmitters on reward-seeking behavior. Int Arch Med 7, 29, (2014).

60 Kim, J. H., Austin, J. D., Tanabe, L., Creekmore, E. & Vezina, P. Activation of group II mGlu receptors blocks the enhanced drug taking induced by previous exposure to amphetamine. Eur J Neurosci 21, 295–300, (2005).

61 Karler, R., Calder, L. D., Chaudhry, I. A. & Turkanis, S. A. Blockade of “reverse tolerance” to cocaine and amphetamine by MK-801. Life Sci 45, 599–606, (1989).

62 Wirtshafter, D. & Stratford, T. R. Evidence for motivational effects elicited by activation of GABA-A or dopamine receptors in the nucleus accumbens shell. Pharmacol Biochem Behav 96, 342–346, (2010).

63 Ranaldi, R. & Poeggel, K. Baclofen decreases methamphetamine self-administration in rats. Neuroreport 13, 1107–1110, (2002).

64 Voigt, R. M., Herrold, A. A., Riddle, J. L. & Napier, T. C. Administration of GABA(B) receptor positive allosteric modulators inhibit the expression of previously established methamphetamine-induced conditioned place preference. Behav Brain Res 216, 419–423, (2011).

65 Snider, S. E., Hendrick, E. S. & Beardsley, P. M. Glial cell modulators attenuate methamphetamine self-administration in the rat. Eur J Pharmacol 701, 124–130, (2013).

66 Worley, M. J., Swanson, A. N., Heinzerling, K. G., Roche, D. J. & Shoptaw, S. Ibudilast attenuates subjective effects of methamphetamine in a placebo-controlled inpatient study. Drug Alcohol Depend 162, 245–250, (2016).

67 Saika, F. et al. Upregulation of CCL7 and CCL2 in reward system mediated through dopamine D1 receptor signaling underlies methamphetamine-induced place preference in mice. Neurosci Lett 665, 33–37, (2018).

68 de Guglielmo, G. et al. Increases in compulsivity, inflammation, and neural injury in HIV transgenic rats with escalated methamphetamine self-administration under extended-access conditions. Brain Res 1726, 146502, (2020).

69 Koob, G. F. & Volkow, N. D. Neurobiology of addiction: a neurocircuitry analysis. Lancet Psychiatry 3, 760–773, (2016).

