## Supplementary Data for "PARKIN REGULATES DRUG TAKING-BEHAVIOR IN RAT MODEL OF METHAMPHETAMINE USE DISORDER"

[b] College of Pharmacy, Natural and Health Sciences
Manchester University
10627 Diebold Rd,
Fort Wayne, IN, USA 46845

[c] Brain Mind Institute
École Polytechnique Fédérale de Lausanne
AI 2240 (Bâtiment AI)
Station 19
CH-1015 Lausanne
Switzerland

Correspondence should be addressed to:
Anna Moszczynska, Ph.D.
Department of Pharmaceutical Sciences
Eugene Applebaum College of Pharmacy and Health Sciences
Wayne State University
259 Mack Ave,
Detroit, MI, USA 48201


**METHODS**

**Stereotaxic Surgery and Parkin Overexpression**

Rats were anesthetized by 4% isoflurane and maintained under anesthesia at 2% isoflurane. Non-coding or *parkin*-coding adeno-associated vector 2/6 (AAV2/6 or AAV2/6-parkin) was bilaterally (behavioral experiments) or unilaterally (PARKIN level assessment) injected at a 16° angle into the nucleus accumbens (NAc) with the following coordinates: +1.8 A/P, ±3.2.M/L, -7.6 D/V from bregma, according to the Paxinos and Watson[1] brain atlas. Viral suspensions were injected with a 5µl Hamilton syringe at a rate of 0.15µL/min using a syringe pump (Harvard Apparatus, Holliston, MA, USA). The syringe was left in place for 5min at −7.6mm, then withdrawn to −6.6mm and −5.5mm, staying in each location for 5min, then raised slowly out of the brain over 5 minutes. The viral suspensions were produced and titrated, as described previously[2] at Swiss Federal Institute of Technology Lausanne (EPFL), Switzerland, and kindly gifted by Dr. Bernard Schneider. The control AAV2 ($2\times10^7$ TUs/side) or AAV2/6-parkin ($2\times10^7$ or $6.5\times10^7$ TUs/side) was microinjected at a volume of 2µL. For sham surgeries, 2µL PBS was microinjected.

**Extended-Access METH Self-Administration (EA METH SA)**

Standard operant-conditioning chambers with two retractable levers for the rat were used (MED-008-CT-B2, Med Associates Inc., St. Albans, VT, USA). D-Methamphetamine HCl (Sigma-Aldrich, St. Louis, MO) or saline (2mg/ml) was delivered through an 18-gauge polyethylene drug delivery tubing via a syringe pump (Med Associates Inc.). During training, active lever presses were reinforced on an FR1 20-s timeout reinforcement schedule and METH infusions were paired with a 5-s tone-light cue. The injected volume was adjusted daily according to the weight of each rat to deliver a METH dose of 0.1mg/kg/injection. Experiments were controlled and recorded by microcomputers using MED Associates interfaces and software. Starting one week before the surgery, and during the whole SA procedure, rats were maintained on an inverse 12h light/dark cycle (lights on at 7:00pm). Rats underwent catheterization with Silastic catheters (C30PU-RJv1303, Instech Lab., Inc., Plymouth Meeting PA, USA) implanted into the right external jugular vein. Following surgery, the IV catheter was flushed daily during the first week with 0.2–0.3mL of sterile saline containing heparin (1.25units/mL) and gentamicin (10mg/mL), followed by 0.2–0.3mL heparin/dextrose solution (SAI Infusion Technologies, Lake Villa, IL, USA). The IV catheters were flushed daily with sterile saline before the start of each session and with the heparin/dextrose solution at the end of each session to maintain catheter patency. All animals were food trained for 2hrs during two consecutive days at fixed-ratio one (FR1) to condition them for lever pressing. A maximum of 20 lever presses was allowed during food training. Animals were rested for one day at the end of food training. METH SA sessions lasted 15hrs a day and were conducted for 10 consecutive days between 6pm and 9am. During the first three days of METH SA, each lever press response (FR1) delivered 0.1mg/kg of METH followed by 20-s timeout period, during which the stimulus light was off and lever in the retracted position. During the next three days of METH SA (days 4-6), FR2 was used whereas during days 7-10, FR5 was used.

**METH Conditioned Place Preference (CPP)**

Place conditioning and preference-testing were conducted in 20×60×20cm plexiglass chambers. One chamber had striped black-and-grey walls and the other solid grey walls. All experiments were recorded by a ceiling-mounted Logitech camera. The total time spent in each compartment was determined by center point detection tracking using EthoVision XT software (Noldus Information Technology, Wageningen,

Netherlands). An experimenter blind to animal condition verified the accuracy of all computer tracking. Place conditioning was performed over ten consecutive days as follows: day 1 – pre-testing phase, days 2-9 – conditioning phase, day 10 – preference-testing phase. On day 1, animals could explore either compartment for 20min to determine if there was a baseline preference towards a specific compartment. METH was paired to the less-preferred compartment. If an animal did not display a preference towards a specific compartment, METH was paired to a compartment by random assignment. Rats were injected (i.p.) with D-methamphetamine HCl (Sigma-Aldrich, St. Louis, MO, USA) (4mg/kg) or saline (1mL/kg) on alternate days and confined to compartments for 30min. On the test day, rats were allowed to explore the compartments for 20min. Change in time spent in the METH-paired compartment was calculated by subtracting the total time spent in the non-preferred side on day 1 from the total time spent in the non-preferred side on day 10.

### RESULTS

#### Body Weight Loss During EA METH SA

The PKO rats weighed more than WT rats but lost weight during METH SA at the same rate as WT controls (Fig.S1A). The PO-NAc rats weighed slightly more than WT controls and lost body weight at the same rate as WTs as well (Fig.S1B). This difference in body weigh was likely due to a small difference in age (several days) between the WT and PO-NAc rats. The PKO and PO-NAc rats did not differ in respect to weight before and during METH SA (Fig.S1C).

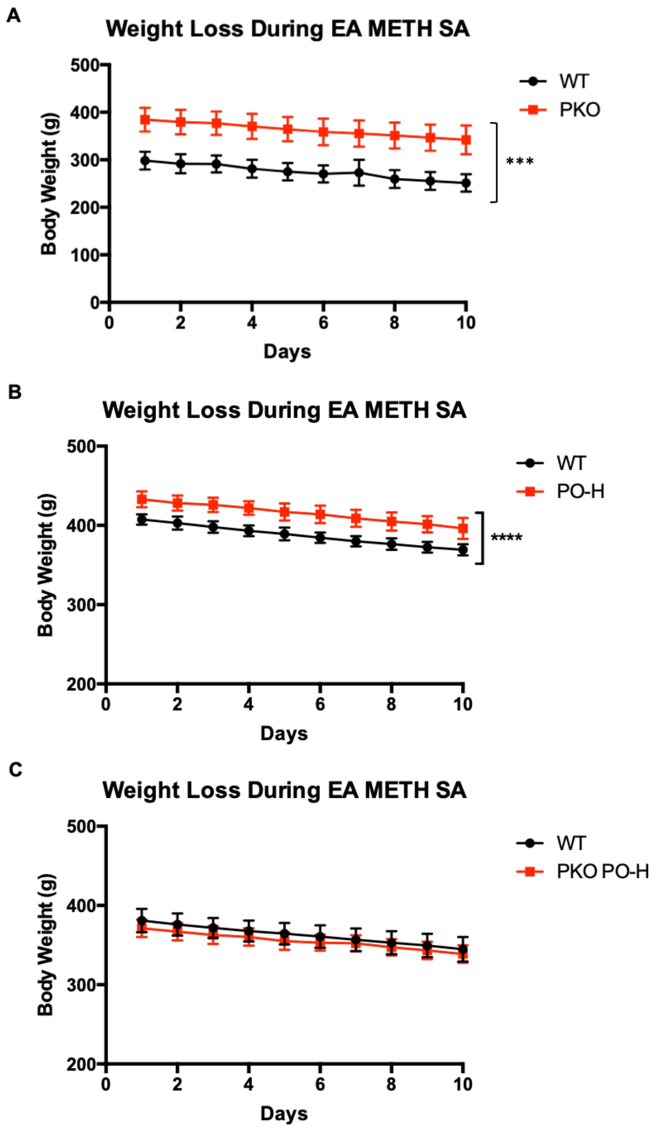


**Figure S1.** Weight Loss During EA METH SA in wild-type (WT), *parkin* knockout (PKO), WT rats overexpressing PARKIN protein in the nucleus accumbens (PO-H), and PKO rats overexpressing PARKIN in the NAc (PKO PO-H). Abbreviations: PO-H, high parkin overexpression (>10-fold).

### REFERENCES

1 Paxinos, G. & Watson, C. The Rat Brain in Stereotaxic Coordinates. *Academic Press*, (2013).

2 Dusonchet, J., Bensadoun, J. C., Schneider, B. L. & Aebischer, P. Targeted overexpression of the parkin substrate Pael-R in the nigrostriatal system of adult rats to model Parkinson's disease. *Neurobiol Dis* **35**, 32-41, (2009).
